# Flexible fitting to infer atomistic-precision models of large-amplitude conformational dynamics in biomolecules from high-speed atomic force microscopy imaging

**DOI:** 10.1101/2025.05.14.653975

**Authors:** Romain Amyot, Osamu Miyashita, Xuan Wu, Kazusa Takeda, Noriyuki Kodera, Hiroki Konno, Florence Tama, Holger Flechsig

## Abstract

High-speed atomic force microscopy (HS-AFM) experiments allow direct observation of biomolecular dynamics at the single-molecule level, acquiring a large amount of topographic imaging data that visualizes changes of the molecular surface during functional activity over an extended period of time. Since images have no atomistic resolution, a major challenge has been to develop post-experimental computational methods to infer atomistic information from measurements. The recently developed NMFF-AFM flexible fitting method provides a computationally efficient approach promising to infer atomistic-precision models of conformational dynamics from resolution-limited AFM imaging data. We report the software integration of this method into the well-established BioAFMviewer platform and demonstrate first applications to experimental HS-AFM imaging data. To facilitate applications, we developed a direct workflow from raw experimental AFM data to visualization and analysis of fitting results. The presented applications to experimental data of a single protein domain, a protein complex, and a megadalton size protein filament demonstrate versatility of NMFF-AFM modelling to reproduce large-amplitude conformational motions of biomolecular dynamics from HS-AFM imaging. As a first step towards automated large-scale analysis of AFM imaging data, we furthermore demonstrate reconstruction of an atomistic molecular movie of protein dynamics, involving large-amplitude conformational transitions, from a measured HS-AFM movie sequence. Implementation of flexible fitting within the standalone user-friendly interactive BioAFMviewer software opens the opportunity for broad range applications to facilitate the understanding of resolution-limited HS-AFM measurements.

## Introduction

Despite the recent revolution in structural biology techniques [1,2] and the use of artificial intelligence for solving the protein structure prediction problems [3,4], revealing the dynamical mechanisms by which proteins implement their biological function remains an outstanding challenge. Well-established single-molecule experimental methods such as fluorescence spectroscopy can make predictions about the dynamical behavior based on optical measurements, but individual molecules themselves are invisible. High-speed atomic force microscopy (HS-AFM) is the only experimental technique to directly observe single proteins during their functional activity under near-physiological conditions [5]. By rapidly scanning a protein placed on a solid substrate with a probing tip, conformational dynamics are monitored as consecutive topographic images visualizing changes in the molecular surface. Directly filming the molecular dynamics of individual proteins their interactions with other biomolecules, and functional assemblies of them, has made important contributions for the understanding of biological processes with interdisciplinary applications in the field of Life Science [6].

HS-AFM can image proteins during their operation over a long time, acquiring a large amount of imaging data, but intrinsic experimental limitations prevent a detailed understanding of the observed dynamics. Visualized are only motions of the biomolecular surface by using a scanning tip that is typically larger by one order of magnitude compared to atomic size, preventing to resolve details at the sub-nanometer range. Although recent technological improvements to speed up scanning [7,8], approaching the time resolution of 10 ms, are promising to expand the range of observable biomolecular dynamics, AFM measurements alone will inevitably provide insufficient information for detailed understanding of biomolecular function.

To fully exploit the explanatory power of HS-AFM measurements, computational methods to reconstruct three-dimensional structures and conformational dynamics with atomistic-resolution from post-experimental analysis of imaging data have been developed in the past years [9]. Such approaches are based on integrating the vast amount of available high-resolution structural data of proteins and molecular modelling methods with HS-

AFM data. Previously developed rigid-body fitting [10,11], correlating known 3D atomistic static structural data with measured topographies, considerably restricts the interpretation of dynamic topography imaging data. On the other side, flexible fitting methods considering conformational changes of structural data have been developed to provide feasible atomistic models of dynamic conformations from imaging data [12-14]. A major drawback in current methods is that dynamically steering an input structure into fitting a target AFM image relies on exhaustive sampling of conformations. Hence, the massive computational expense required to conduct flexible fitting has so far prevented applications for the analysis of AFM measurement data.

Recently, two separate studies have proposed highly accelerated flexible fitting methods, NMFF-AFM (normal mode flexible fitting AFM) [15] and AFMfit [16]. For the computational sampling of biomolecular dynamics to identify an atomistic structure that matches an experimental image, these methods focus on the reduced conformational subspace derived from normal mode calculations of an elastic-network presentation of a known biomolecular atomistic structure. While applications of AFMfit are missing to resolve motions that are significantly different from the initial structure, the NMFF-AFM method has demonstrated the potential of flexible fitting to reproduce atomistic-precision models of large-amplitude conformational transitions observed under HS-AFM microscopy [15].

Here, we report the software implementation of NMFF-AFM method into the well-established standalone BioAFMviewer platform [17], allowing for convenient applications to experimental imaging data within a user-friendly interactive graphical interface with no dependencies on other software. While in the original publication [15] the model performance was verified in control applications of synthetic AFM data, we now present the developed software-integrated workflow of flexible fitting and demonstrate several applications to HS-AFM imaging data.

As we show, flexible fitting achieves prediction for atomistic models of large-amplitude conformational motions from HS-AFM images with remarkable computational efficiency, requiring as few as tens of seconds calculation time for the presented applications to a single protein domain and a protein complex. This significant advancement of computational modelling opens the opportunity for automatized large-scale analysis of AFM imaging data, also including the interpretation of measurements for large-size protein aggregates such as filaments.

To outline such prospects, we demonstrate application of the NMFF-AFM method to reconstruct a molecular movie of atomistic-precision models of protein dynamics involving functional conformational transitions from a HS-AFM movie. The computational efficiency is furthermore highlighted in the application of flexible fitting for a megadalton size actin filament to provide an atomistic model from topographic imaging, a task which would require surreal computational expense if attempted by fitting using molecular dynamics simulations.

Integrating flexible fitting into a user-friendly software workflow – from raw experimental AFM imaging data to visualization and analysis – is expected to enable a broad range of applications by the Bio-AFM community. This will facilitate a detailed understanding of molecular processes at the nanoscale from topographic AFM measurements.

## Software integration of flexible fitting

We first summarize the concept of the NMFF-AFM computational method and refer to the original publication [15] for a detailed description. This method relies on an iterative modeling scheme to gradually deform an atomistic biomolecular structure towards a dynamic conformation that fits into a target experimental AFM topographic image. The deformations are generated by computing conformational motions arising from the normal mode analysis of an elastic network representation of the biomolecular structure.

Starting with an initial structure, known either from previous experimental data (Protein Data Bank structures) or AlphaFold predictions, the iteration scheme performs, in each step, first normal mode analysis to generate a set of slightly deformed structures representing motions of individual modes and amplitudes. Next, AFM imaging simulation to translate this set of atomistic conformations into pseudo-AFM topographic images allows comparison with the target AFM image to select a single normal mode motion and its direction (sign of the amplitude) that best improves image similarity. The deformed structure obtained by using the selected motion then enters the next round of iteration (with its elastic network possibly changed). The iteration scheme terminates when fitting deformation cannot much further improve the image similarity of the simulated AFM topography for the deformed structure and the experimental AFM image.

The method has been validated in applications of different proteins with various characteristics of conformational changes under idealized conditions [15], where the target AFM image of a protein was generated as a simulated topography from a known atomistic structure (ground truth). Flexible fitting starting from another known structure of the same protein in a completely different conformation was then able to sufficiently well reproduce the ground truth structure.

We implemented the NMFF-AFM method within the BioAFMviewer software for convenient applications to *real-world* experimental HS-AFM imaging data. The BioAFMviewer platform was designed towards integrating all available high-resolution structural and modeling data of biomolecules with resolution-limited scanning probe measurement data. The software package contains a high-quality molecular viewer for the 3D visualization of biomolecular structures combined with synchronized computation and visualization of corresponding simulated AFM images (live simulation AFM), allowing to correlate atomistic structures with measured topographies as previously demonstrated by applications of the integrated rigid-body fitting method [11].

With the prospects of flexible fitting allowing for efficient applications to large sets of HS-AFM imaging data, we have also implemented a new software tool for automated integration of experimental AFM raw data in the BioAFMviewer, the BioAFM-ASD viewer. This app processes data encoded in the ASD file format (a standard format used by AFM experimentalists) to visualize the dynamics of measured biomolecular surface topographies as 2D color-coded multi-frame images, thus establishing a communication pipeline to correlate experimental imaging data with simulated AFM topographies of dynamic atomistic conformations generated during fitting. The BioAFM-ASD viewer includes options for post-experimental image processing prior to the application of flexible fitting, such as AFM-stage tilt correction, Gaussian filtering of pixel information along the scanning grid, and a method for automated molecule recognition to define the target AFM image relevant for fitting (see Methods for details).

For the convenient integration of the NMFF-AFM workflow we have implemented a flexible toolbox app. Like existing BioAFMviewer apps, it provides a user-friendly interface to choose parameters of the model and the fitting process. The app directly communicates with the BioAFMviewer molecular viewer, simulation AFM, and the ASD-viewer, to conduct the fitting procedure. Thereby, it integrates the cycle of iterative normal mode analysis of the biomolecular structure and comparison of simulated topographies for dynamic conformations with the target experimental AFM image towards eventually identifying the best matching fitting result. After completion of the fitting process, the app allows for direct visualization and analysis of obtained fitting results. We refer to the Methods section for further detailed information.

Fig. 1 shows a schematic workflow for the application of the NMFF-AFM method within the BioAFMviewer, involving the three stages of loading data, application of the flexible fitting toolbox, and graphical output combined with data analysis.

**Fig. 1.**
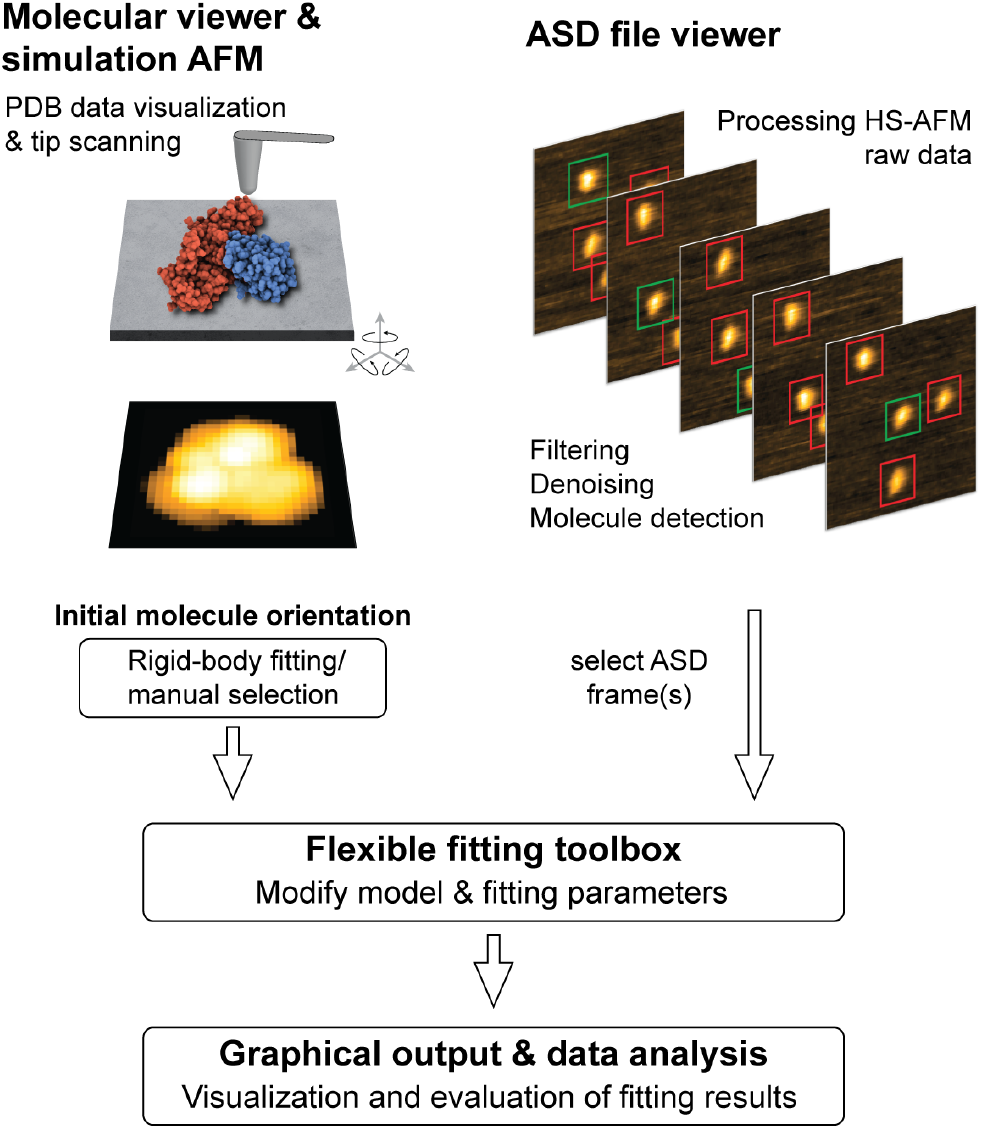
Workflow for BioAFMviewer software integration of NMFF-AFM flexible fitting to AFM imaging data. The BioAFMviewer molecular viewer allows 3D visualization of available biomolecular structures combined with interactive live simulation AFM to compute and visualize corresponding pseudo-AFM topographic images to be correlated with experimental images during fitting processes. The developed ASD file viewer incorporates the import of AFM raw data to visualize measured topographic images and provides image processing methods including molecule recognition to autonomously identify targets for fitting. The flexible fitting toolbox app allows convenient selection of model and fitting parameters and integrates all computations of the NMFF-AFM iteration scheme. The interactive BioAFM software interface allows visualization of fitting results, data analysis, and convenient export of all obtained data.

## Applications to HS-AFM imaging data

### HECT domain large-amplitude conformational transition

The first application is for HS-AFM imaging of the structural dynamics of HECT (homologous to E6AP C-terminus) domain (∼350 amino acids) that plays an important role in post-translation protein ubiquitination by HECT E3 ligase enzymes to regulate numerous cellular processes [18]. HS-AFM experiments visualized highly dynamic topographic changes showing frequent transitions between spherical and oval molecular shapes (Ref. [19], and Fig. 2c for a few examples), which suggests large-amplitude conformational motions within the HECT domain under physiological conditions. Previous application of rigid-body fitting allowed to assign known static atomistic structures in very different conformations to either of the measured topographic shapes [19].

**Fig. 2.**
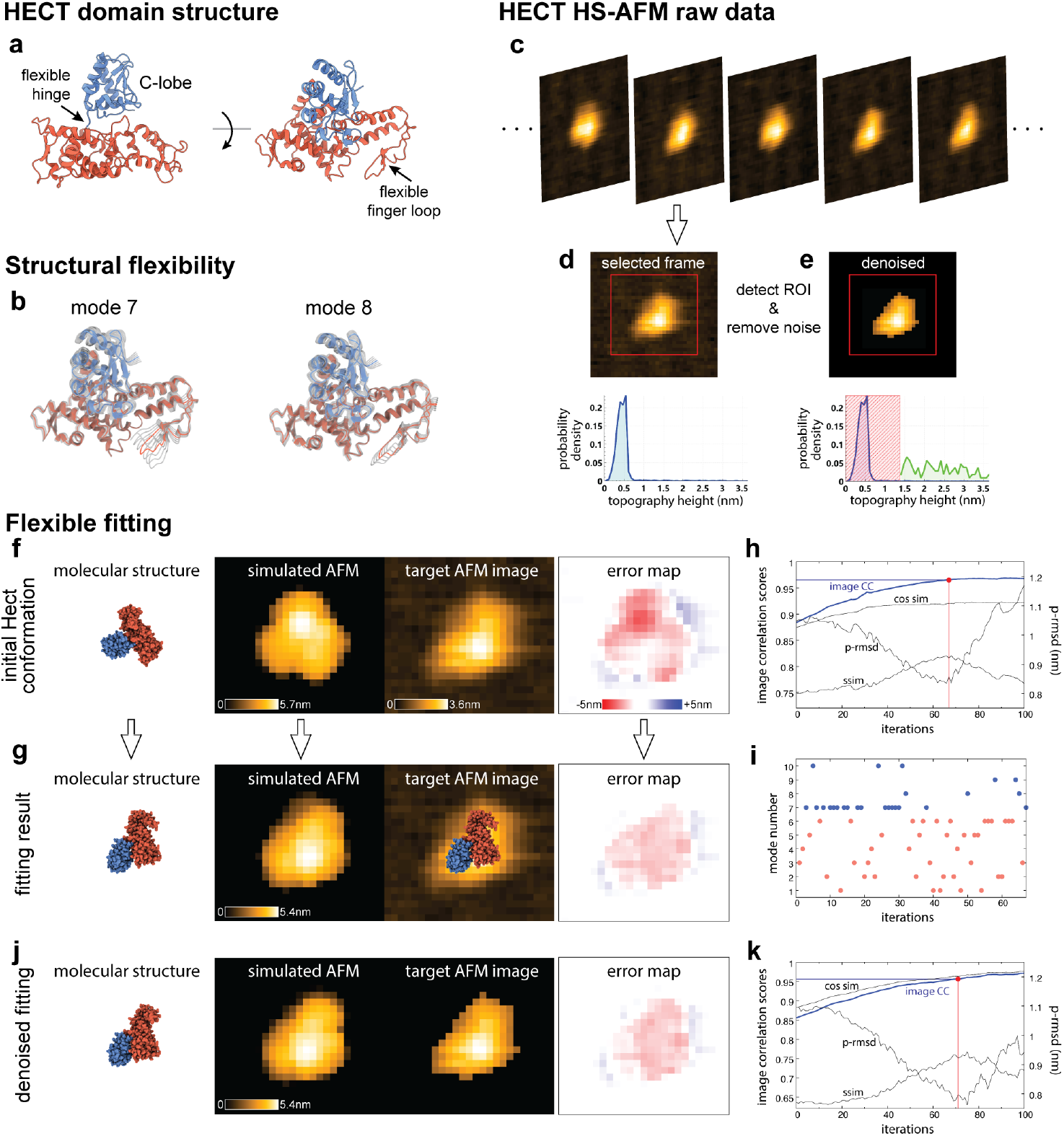
HECT domain large-amplitude conformational transition. **(a)** Ribbon representation of HECT domain structure in the catalytic conformation. The C-lobe domain (blue) and finger loop region are indicated. **(b)** Snapshots of structure changes in normal modes 7 and 8, respectively. The HECT structure is colored and deformed conformations are shown in transparent grey. **(c)** Examples of typical HS-AFM images for HECT topographies in spherical and oval shapes. **(d)** The ASD image selected for fitting and the probability density of topography heights. **(e)** The same ASD image after applying filtering according to the height threshold determined by the Otsu method (denoised image). The corresponding height probability density shows variations of the HECT domain shape by green color, rescaled after erasing the background (shown as a red colored box for illustrations purpose only). In (d,e) the region of interest (ROI) for fitting is shown in red. **(f)** Side-by-side image presentation of the initial HECT conformation for fitting (atomistic VdW representation), the corresponding simulated AFM image, the ROI of the target AFM image, and the error map. **(g)** The corresponding images after application of flexible fitting. **(h)** Plots of image similarity scores along the fitting process. The graph of the image cc score used for fitting is shown in blue; graphs of other scores are displayed in grey (for the p-rmsd an alternative scale is used). The iteration point determined by the fitting termination criterion is marked as a red dot. **(i)** Plot of the mode number selected along the fitting process; deformation modes are shown in blue. **(j)** Fitting results employing the denoised target AFM image (representation similar to that in panel (g)). **(k)** Corresponding plots of image similarity scores.

Here, we demonstrate flexible fitting to model large-amplitude transitions in applications for HECT domain HS-AFM imaging data. Fig. 2a shows the HECT domain structure in its so-called catalytic conformation (PDB ID 3jvz) which has been identified to represent measured spherical shape AFM topographies [19]. The conformation is characterized by a globular C-lobe domain connected in a central position with respect to the rest of the protein via an unstructured linker. Moreover, an extended finger loop is present in another protein domain. Previous analysis of crystallographic B-factors and molecular dynamics simulations revealed structural flexibility of those protein parts [19]. Normal mode analysis (NMA) of this HECT structure preceding flexible fitting shows that the first four deformation modes indeed describe relative motions of the C-lobe domain and the finger loop; see Fig. 2b for snapshots of structural changes in modes 7 and 8, respectively.

In our setup of flexible fitting, the HECT catalytic conformation is the starting structure. Fig. 2c shows typical HS-AFM images visualizing spherical and oval molecular shapes. As a target experimental image for fitting, we selected an image visualizing the HECT domain in the oval-shaped topography, which was previously correlated with a static structure in a completely different HECT conformation by rigid-body fitting [19]. For the selected ASD frame (Fig. 2d) we first computed the probability density of topography heights, which shows a narrow distribution (peak at ∼0.5 nm) corresponding to imaging noise of the AFM mica substrate (Fig. 2d). Application of the Otsu algorithm [20] determined a height threshold of ∼1.4 nm to separate the HECT shape from the noisy background (denoised image shown in Fig. 2e) and to determine the region of interest (ROI) for fitting (shown in Fig. 2d,2e). We performed flexible fitting for both noisy and denoised experimental target images.

To choose the initial molecular orientation for flexible fitting, we use information from previous rigid-body fitting of the HECT structure to HS-AFM images [19] (see Methods). Fig. 2f shows the initial molecular orientation, the corresponding spherical shape simulated AFM topography, the target AFM image in the oval shape topography, and the error map visualizing their deviations. The discrepancies between the simulated and experimental AFM image before fitting are characterized by differences in the overall topography shape (spherical versus oval) and in the pattern of height protrusions.

The results obtained from flexible fitting (see Methods for details on parameters) are shown in Fig. 2g. During the fitting process, significant structural changes of the HECT domain are observed, involving internal large amplitude conformational motions of the flexible C-lobe domain and the finger loop, combined with rigid-body rearrangements (see SI Movie 1). Changes between the initial structure and that obtained after fitting are quantified by an RMSD value of 14.5 Å, and 8.4 Å after structure alignment. The simulated AFM image of the fitted structure nicely resembles the oval shape and protrusions of the experimental AFM image, as evidenced by the corresponding error map. The fitting process is visualized in a side-by-side view of the molecular movie, the simulation AFM movie, and the target AFM image, and as a superposition of the molecular movie with the target AFM image (SI Movie 1).

The quantitative improvement in image similarity scores along the fitting process is shown in Fig. 2h. The image c.c. score used to drive the fitting process changes from 0.88 for the initial conformation and saturates at a value of 0.97, where the termination criterion detected the iteration step to determine the structure considered as the fitting result (see Methods). This choice is consolidated by monitoring other image similarity measures during fitting (p-rmsd, ssim; Fig 2h), showing that further conformational changes are unfavorable to improve image similarity. The distribution of selected normal modes along the fitting process (Fig. 2i) confirms that motions of the HECT domain result from a combination of rigid-body rearrangements and internal conformational changes which are dominated by the softest deformation mode.

We repeated the exact same flexible fitting using the denoised AFM image (Fig. 2e) as a target. The results are shown in Fig. 2j,k, and the fitting process is visualized in SI Movie 2. Comparing the molecular structures obtained from the two fittings (Fig. 2g,j) shows some differences, most visible in the position of the flexible finger loop region. The overall differences are quantified by a RMSD value of 4.4 Å (without alignment). However, the two obtained structures align quite well by a much smaller RMSD of 1.9 Å. The implication of this observation is that differences between the fitting results are mainly attributed to a change in the orientation of the HECT structure relative to the AFM substrate, while similar internal large amplitude conformational motions towards the target AFM image are produced during flexible fitting (regardless of the presence of noise). We argue that for the denoised case the fitting procedure generated additional rotations of the HECT structure (see SI Movie 2) to improve image similarity toward the target AFM image in which denoising has removed the boundary of the protein topography at heights comparable to the background noise.

The quantitative improvement in image similarity scores along the fitting process (see Fig. 2k) shows a change in the image c.c. from 0.86 to 0.96 (at the termination iteration) comparable to that for fitting to the noisy AFM image. Note, that for the denoised fitting the cosine similarity score which does not consider the average of topography information and the image c.c. are similar.

To compute simulated AFM images during fitting, we had to decide on parameters characterizing the tip shape geometry by its probe sphere radius and the cone half angle. In particular, the size of the probe sphere affects simulated topographies [10,17], thereby possibly changing the results of flexible fitting to the target experimental AFM image. We addressed this issue by performing flexible fitting for different tip shapes varying in the size of the probe sphere radius (R=1nm, R=2nm, R=3nm). All three fitting procedures generated similar conformational changes of the flexible C-lobe domain and the finger loop (see SI Fig. 1, and SI Movie 3), and showed comparable improvements in the image similarity score. For the fitting presented above, we favored fitting using a tip shape with a probe sphere radius of 2nm and a cone half angle of 10 deg, recording the highest image c.c. between the simulated AFM image of the fitted model conformation with the AFM target image.

Finally, we aimed to compare the dynamic HECT domain conformation obtained from flexible fitting with the static structure in the L-shape conformation (PDB ID 1D5F) that was previously correlated with measured oval shape HS-AFM topographies based on rigid-body fitting [19] and is considered functional to transfer ubiquitin to target proteins [21]. Since the structures represent different types of HECT domains (E6AP-type HECT used for flexible fitting and the protein used in experiments; NEDD4L-type HECT for L-shape) with poor sequence similarity, we characterized the conformations in terms of their domain geometry rather than using atomistic RMSD analysis. The position and orientation of the C-lobe domain relative to the N-lobe domain was characterized by the distance between their geometric centers and a hinge angle (see Methods), showing very different values for the HECT domain catalytic conformation used as the initial state for flexible fitting (angle=108 degrees; D=3nm) and the static structure L-shape conformation (angle=67 degrees; D=3.1nm) (see SI Fig 2). As we find, the HECT protein conformation obtained from flexible fitting is similar to the L-shape geometry (angle=79 degrees; D=3nm), thus supporting experimental observations of E6AP-type HECT to adopt functionally important conformations under equilibrium physiological conditions.

### HECT domain complex with bound E2 enzyme

To demonstrate the versatility of integrated fitting toolbox, we considered as a second example the application to a protein complex, combining rigid-body fitting to determine an initial molecular orientation and optimal tip-shape parameters with subsequent flexible fitting to provide an atomistic model of experimental HS-AFM imaging.

Before the HECT-E3 ligase can ubiquitinate target proteins, it must accept a ubiquitin (Ub) molecule delivered by the small E2 enzyme in a first step. HS-AFM experiments including both HECT and E2-Ub enzymes could visualize topographies that were different from spherical or oval shapes of a single HECT domain [19], but it was not previously possible to clearly verify the formation of a protein complex using rigid-body fitting of available static atomistic structures. We therefore applied a combination of rigid-body and flexible fitting to provide an atomistic model for the measured HS-AFM image.

Fig. 3a shows the results of rigid-body fitting of the HECT-E2 protein complex (494 amino acids, PDB ID 1c4z) obtained by the fitting toolbox in the BioAFMviewer software [11]. The simulated AFM image of the molecular orientation compares to the HS-AFM image by an image c.c. score of 0.94. Indeed, the agreement is good for the region corresponding to the topography of the HECT domain. Notably, the predicted orientation of HECT is very similar to those obtained from previous rigid-body fitting of the single HECT domain to several measured HS-AFM images [19]. However, the agreement between simulated and HS-AFM image is less good for the part that represents the bound E2 enzyme, as indicated by the error map. This shows that the observed dynamic HECT-E2 complex is clearly different from that of the crystal structure. Before proceeding with flexible fitting, we used the Quick-Fit function of the BioAFMviewer which refines the result of rigid-body fitting by iteratively sampling orientations that further improve image similarity towards the target AFM image considering different tip shape geometries to compute simulated AFM images. Collecting such results, we could therefore construct a landscape of image c.c. values for varying tip shape parameters shown in Fig. 3b. While the cone angle value does not affect the comparison of simulated images with the experimental AFM image, a strong dependence on the probe sphere radius can be clearly seen. Deviations from R=2.0 (used in the previous example of flexible fitting) to either sharper or more blunt tips result in decreasing image c.c. scores. The optimal tip shape parameters are determined as (*R* = 3*nm*, ϑ = 14°).

**Fig. 3.**
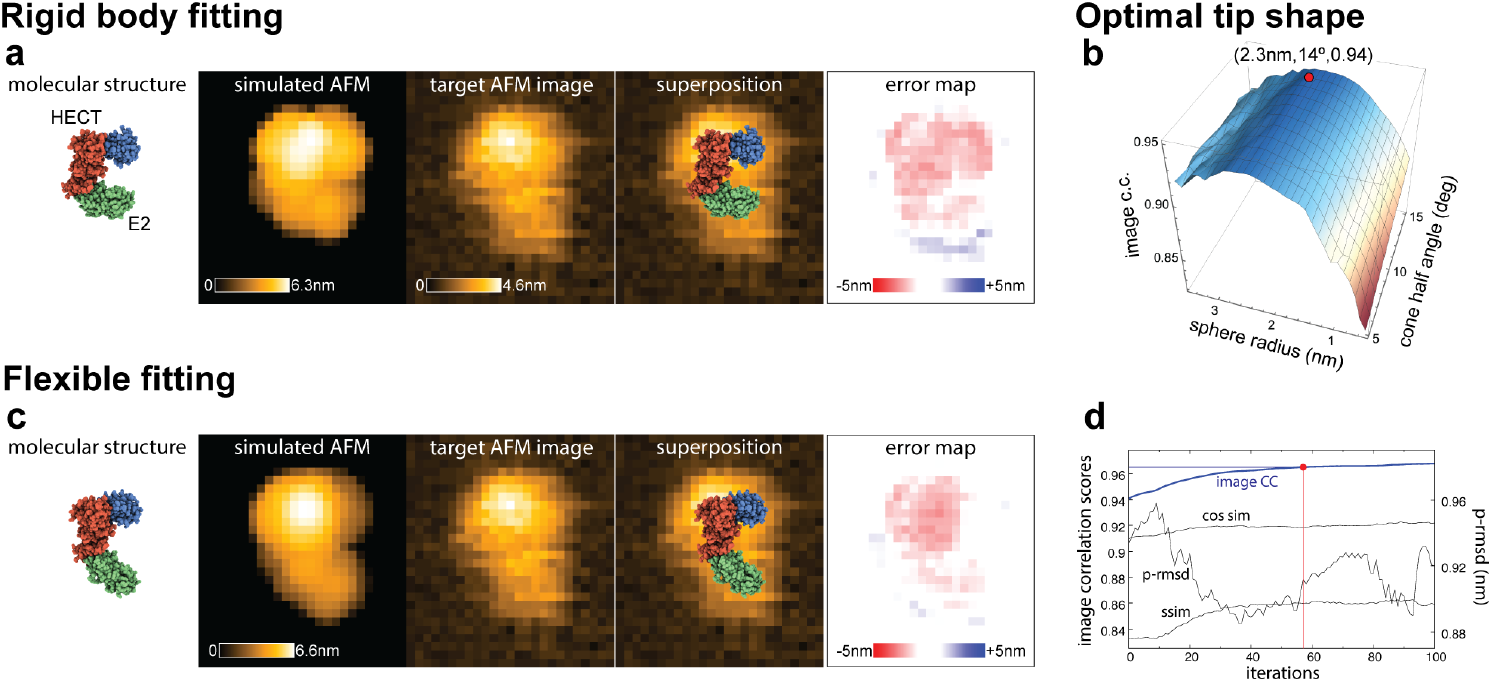
HECT protein complex with bound E2 enzyme. **(a)** Side-by-side view of the HECT-E2 structure in the orientation obtained from rigid body fitting, the corresponding simulated AFM topography, the target experimental AFM image, the superposition of the molecular structure with the AFM image, and the error map between simulated and target AFM images. **(b)** Landscape of image c.c. values scoring similarity of simulated AFM images with variable tip shape parameters and the target AFM image obtained from the quick fit algorithm of refined rigid body fitting. The optimal choice is indicated by a red dot with the parameter and score values shown. **(c)** Results from flexible fitting in the same representation as shown in panel (a). **(d)** Plots of image similarity scores along the fitting process. The graph of the image c.c. score used for fitting is shown in blue; graphs of other scores are displayed in grey (for the p-rmsd an alternative scale is used). The iteration point determined by the fitting termination criterion is marked as a red dot.

For the application of flexible fitting, the HECT-E2 complex is described by a single elastic network, modelling intra and intermolecular interactions in the same way. We used the optimal tip shape for the computation of simulated AFM images and started fitting from the refined molecular orientation of the HECT-E2 crystal structure obtained by rigid-body Quick-Fit. The results from flexible fitting are shown in Fig. 3c (see Methods for details on parameters). The fitting process mainly involves large amplitude motions of the E2 protein relative to the HECT domain quantified by an RMSD value of 10.1 Å and 8.3 Å after structure alignment. The visualization is provided in a side-by-side view of the molecular movie, the simulation AFM movie, and the target AFM image, and as a superposition of the molecular movie with the target AFM image (SI Movie 4). The simulated AFM image of the fitted structure compares well with the target AFM image (image c.c. value of 0.96) both in the overall shape topography and position of protrusions arising from the HECT domain as demonstrated visually by the error map. The quantitative improvement in image similarity scores along the fitting process (Fig. 3d) shows that beyond the iteration step determined by the fitting determination criterion, the image c.c. score used for fitting saturates, indicating no further improvement.

### Reconstructing atomistic precision molecular movies from HS-AFM imaging

The presented applications demonstrate that NMA flexible fitting can capture large-amplitude structural changes by efficiently steering a known static protein structure towards a dynamic conformation representing a plausible atomistic model for a single HS-AFM topographic image. Within the BioAFMviewer software, the performed fitting computations proceeded within a few tens of seconds on normal laptop machines (see Methods section for benchmarking details). This remarkable computational efficiency combined with implementation of automated molecule detection from HS-AFM raw data allows us to extend the application of flexible fitting to reconstruct atomistic conformations from a time series of measured protein topographies.

To provide a demonstration we analyze again HS-AFM data for the structural dynamics of the HECT domain visualizing conformations in the spherical and oval topography shapes, and, transitions between them. We consider experimental data for a single molecule whose dynamic topography is traced in our ASD file viewer to determine for each frame the region of interest which defines the corresponding target AFM image (see Methods for the data set). For each target AFM image, flexible fitting was performed individually, always starting from the same molecular orientation of the HECT domain catalytic conformation used for previous fitting (see Fig. 2f). The model and fitting parameters were also the same, and similar fitting termination criteria were employed to determine the atomistic model behind each target AFM image (see Methods).

Collecting the results of fitting processes, we first quantified the structural change of each obtained conformation compared to the initial structure. We find that conformations group into those with minor changes (RMSD 2-4Å), moderate changes (RMSD 4-7Å), and large internal motions (RMSD > 8Å) (Fig. 4a). HECT conformations obtained from fitting with minor changes are representative of dynamic structures in the near catalytic state (i.e., close to the initial conformation), corresponding to oval shape topographies of the analyzed HS-AFM video frames. Figure 4b shows an example case of such fittings. On the other side, dynamic structures obtained by larger internal motions are representative of conformations corresponding to AFM topographies in spherical-to-oval intermediate shape (see Fig. 4c for an example case), or those which correspond to clearly oval shape AFM topographies (see Fig. 4d for an example case). It should be noted that the result shown in Fig. 4d are consistent with our previous flexible fitting to an AFM image in a different oval shape topography (see Fig. 2g).

**Fig. 4.**
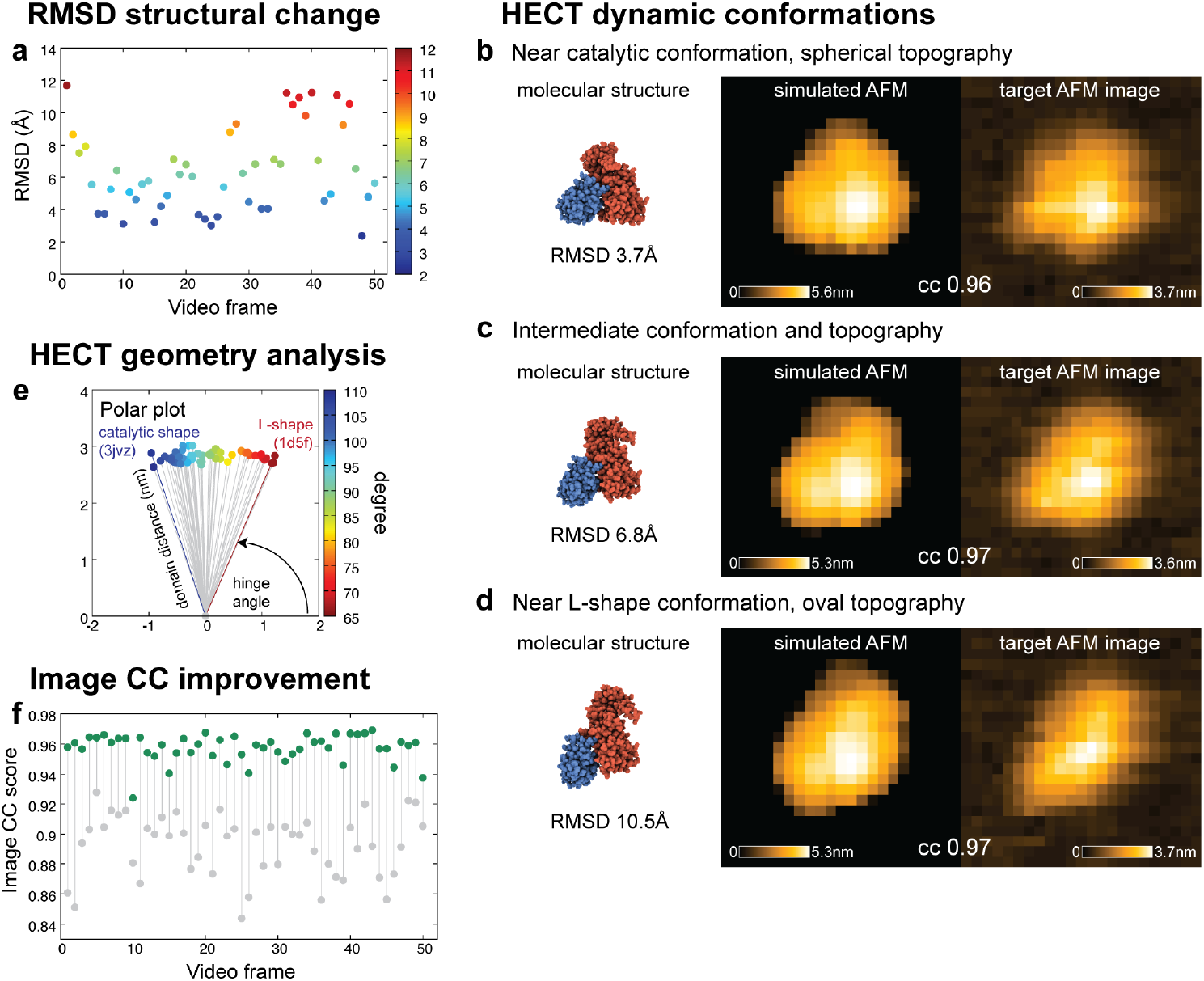
Reconstructing atomistic precision molecular movies from HS-AFM imaging. **(a)** Structure change of dynamic HECT domain conformations with respect to the initial structure as obtained from flexible fitting towards respective video frame target images. The RMSD value representing only internal structural changes is plotted and visualized by a color gradient. Exemplary snapshots representative of dynamic HECT structures in near-catalytic conformation **(b)**, intermediate conformation **(c)**, and near L-shape conformation **(c)**, are shown in a side-by-side view of molecular structure, simulated AFM, and target AFM image. Corresponding RMSD values and image cc values are given. **(e)** Geometry analysis of HECT dynamic conformations from fitting visualized in a polar plot of the C-N-lobe domain distance and the hinge angle (highlighted by color-coded value). Corresponding values for the static catalytic and L-shape PDB structures are indicated. **(f)** Plot of the image cc value for each individual fitting process, visualizing improvement in the image similarity score from values correlating the initial structure simulation AFM image with respective video frame target images (grey colored dots) to correlations obtained after flexible fitting (green colored dots).

In SI Movie 5 the obtained entire HECT dynamics molecular movie is shown in a side-by-side view with the corresponding sequence of simulation AFM images, the sequence of target HS-AFM images, and a superposition of the molecular movie with the HS-AFM movie. The inferred atomistic models for HECT dynamic conformations are characterized by large-amplitude rotations of the flexible C-lobe domain and relative changes of the small finger loop region compared to the HECT catalytic state structure.

The geometry analysis of all collected dynamic HECT conformations indeed shows large variations in the C-N-lobe hinge angle, with values ranging between those for the static catalytic and L-shape structures (Fig. 4e). Here, we recall our previously published findings [19]: First demonstrating by rigid-body fitting that the static catalytic HECT and L-shape conformations can be correlated with spherical shape and oval shape AFM topographies, respectively. Second, from modeling the conformational dynamics along the transition between both states combined with simulation AFM, we previously concluded that spherical-to-oval shape changes in AFM topographies are likely related to the rotation of the C-lobe domain relative to the N-lobe, though it was not possible to directly correlate modeling results with HS-AFM imaging data. The results from flexible fitting, now providing atomistic models for HECT dynamic conformations directly from measured AFM dynamic topographies, allow the detailed level verification for the interpretation of HECT protein functional dynamics from imaging data.

Looking further into the obtained results, we find that in the majority of cases, fitting generated dynamic HECT conformations which can be viewed as high confidence atomistic models of corresponding HS-AFM movie frames, as evidenced by image cc values between simulated AFM images and experimental topographies as high as 0.95 to 0.97 (Fig. 4f). For the few cases with lower image cc values (i.e., 0.92 and ∼0.94) we inspected the individual fitting processes and find two different scenarios. In one scenario, the image cc score quickly increases during a first phase of fitting, then saturates for a few iteration steps, before further increasing almost linearly until the preset last iteration step is reached (see Fig. S3a-c). This rare type of behavior is not well represented when applying our automated fitting termination criteria which as a simplification assumes exponential increase in the image cc improvement as fitting progresses (see Methods), hence resulting in premature termination close to the saturation stage. In such rare cases we repeated the individual fitting procedures, this time extending the number of iteration steps to 200, which allowed saturation of image cc scores to obtain higher confidence atomistic models (see Fig. S3a-c).

In the other scenario, the improvement of image cc score during fitting clearly saturates within the preset range of iteration steps as detected by the automated termination criteria. However, further improvement in image similarity by fitting is hampered due to the presence of experimental artifacts in the corresponding HS-AFM images (see Fig. S3d-f). Such topographies clearly show distortions along the scanning directions indicative of scanning with a contaminated tip or tip parachuting effects, thus limiting the flexible fitting prediction quality.

### Scalability of flexible fitting – application to 4 megadalton actin filament

The computational efficiency of software-integrated NMFF-AFM method shall allow to expand applications to the analysis of AFM imaging data for proteins assemblies, such as filaments with the size of several hundredth of nm.

We provide a demonstration for the scalability of flexible fitting in an application for HS-AFM imaging of the actin filament, which plays an important role for dynamics of the cell cytoskeleton and contraction of muscle cells. The HS-AFM technique was previously employed to observe dynamical changes in the actin filament in response to interactions with binding proteins [22,23], but data analysis could be performed only at the level of measured limited-resolution topographic data. It is therefore generally desirable to infer atomistic models of dynamic actin conformations from AFM imaging for detailed analysis and understanding of measurements.

Here, we provide an example application of flexible fitting considering a single HS-AFM experimental image of the actin filament which was observed in a bent shape (Fig. 5a), much different from the textbook perception being straight. To perform fitting, we first constructed an atomistic model of a straight actin filament consisting of 96 protomer units shown in Fig. 5a (278,304 atoms in total, see Methods). Considering that AFM observations proceed with the actin filament placed on the 2D substrate allowed us to determine a suitable initial orientation of the atomistic model (see Methods for details). Furthermore, the comparison of simulated AFM topographies with the measured HS-AFM image allowed us to estimate tip-shape parameters for fitting as (*R* = 6.5*nm*, ϑ = 10°) (see Methods and Fig. S4).

**Fig. 5.**
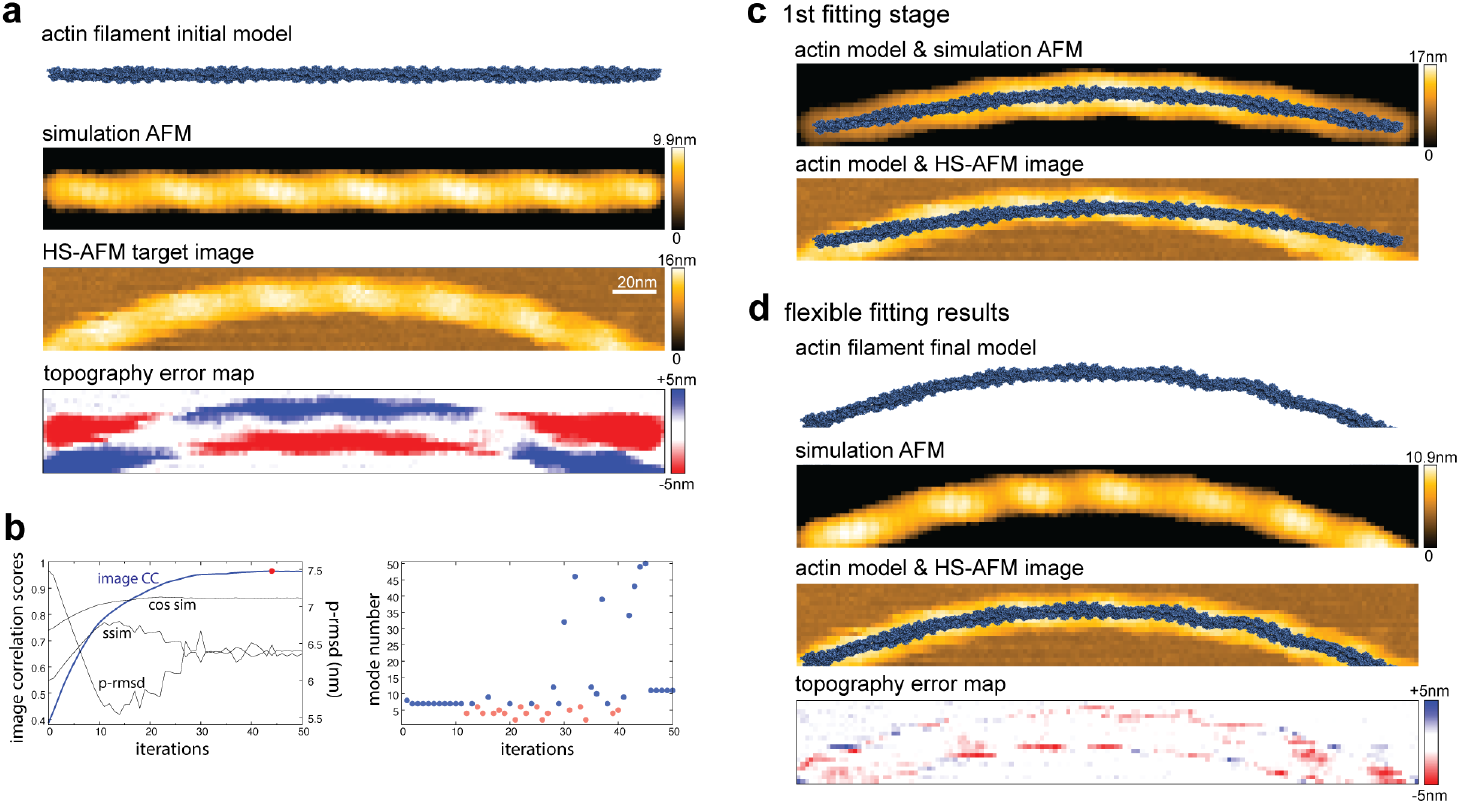
Flexible fitting of 4 megadalton actin filament. **(a)** Initial 96-protomer atomistic model of the actin filament, its corresponding simulation AFM image obtained with optimal tip-shape scanning, the target HS-AFM image of bent actin shape, and the topography error map before fitting (from top to bottom). **(b)** Left: Plots of image similarity scores along the fitting process. The graph of the image c.c. score used for fitting is shown in blue; graphs of other scores are displayed in grey (for the p-rmsd an alternative scale is used). The iteration point to terminate fitting is marked as a red dot. Right: Plot of the mode number selected along the fitting process. Deformation modes are indicated as blue dots. Modes corresponding to rigid-body rotation (number 1 to 3) and translation (number 4 to 6) are shown as red dots. **(c)** Mid-stage result of fitting resulting solely from global bending motions of the actin model favored in the first stage of fitting. The instantaneous actin model at iteration step 11 is shown in superposition with its corresponding simulation AFM image (top), and the target HS-AFM image (bottom). **(d)** Results after finalized flexible fitting. From top to bottom: atomistic model of the fitted actin filament, its corresponding simulation AFM image, its superposition with the HS-AFM target image, and the topography error map.

The results from flexible fitting are shown in Fig. 5 (b-d) (see Methods for parameter details). We find that fitting proceeds in two stages. During the first stage, the fitting algorithm is solely driven to level the differences between the simulated AFM straight topography shape with respect to the measured bent shape topography, since only global bending motions of the actin filament characterized by the first non-trivial normal mode (mode number 7) are performed (see Fig. 5b). During this part of fitting the image cc value already significantly improved from 0.39 to 0.79 (see Fig. 5b). The dynamic atomistic actin filament model resulting from the first fitting stage is shown in Fig. 5c together with the corresponding simulation AFM image and the experimental target AFM image. An interesting observation at this fitting stage is that while globally bent shapes of simulated and measured AFM topographies are already comparable by an image cc value of 0.79, they mismatch in terms of protrusion produced by the periodic arrangement of protomers along the filament. This is because the first stage fitting process also favored global bending motions which deviate the actin filament orientation from the planar AFM x-y surface (see topography height scale of the simulated AFM image in Fig. 5c), apparently to better match the protrusion pattern in the central HS-AFM image region where the actin filament is rather straight.

The second fitting stage is then dominated by rigid-body motions of the actin filament combined with further internal structural changes (see Fig. 5b for the selected normal modes). As we find, the rigid-body rearrangements consist in rotations and translations to place the bent filament back towards the planar orientation in which the simulated AFM image represents well the periodic protrusion pattern of the measured AFM image. Regarding internal structural changes. it is interesting that this fitting stage includes several steps which employ motions of higher-frequency normal modes (numbers 32 to 50). In contrast to low-frequency modes describing homogenous bending of the actin filament, such modes rather correspond to deformations changing locally the arrangement between neighboring protomers, thus generating slight deviations from the global symmetrically bent filament structure. This can be seen in the atomistic model of the actin filament structure after finalized fitting shown in Fig. 5d together with the simulation AFM image, the superposition of atomistic model with the target HS-AFM image, and the topography error map. The confidence of the obtained atomistic model to represent the experimental HS-AFM image is quantified by an image similarity cc score of 0.96 (see Fig. 5b).

The entire fitting process is visualized in a top-to-bottom view of the molecular movie, the simulation AFM movie, the target AFM image, and as a superposition of the molecular movie with the target AFM image (SI Movie 6).

## Summary and Discussion

We provide the software implementation of the flexible fitting NMFF-AFM computational method within the popular BioAFMviewer platform and demonstrate first applications to experimental HS-AFM imaging data. The developed BioAFM-ASD viewer app provides the interface to visualize and analyze measured AFM topographic data, while the NMFF-AFM app provides a graphical interface for flexible fitting and integrates all required computations. The implementation capitalizes on the BioAFMviewer stand-alone high-quality molecular viewer to visualize atomistic biomolecular structures, its synchronously operating AFM simulator, previously implemented rigid-body fitting methods, AFM topography analysis tools, and the graphical interface to directly visualize obtained results and export data. Hence, the established user-friendly environment, with no dependencies on other software products, offers the opportunity for broad applications of flexible fitting within the Bio-AFM community to infer 3D atomistic-precision models of resolution-limited topographic imaging.

The presented applications to experimental observations of a single protein domain, a protein complex, and a megadalton size filamentous protein clearly demonstrate the versatility of flexible fitting modelling large-amplitude conformational transitions to provide atomistic-precision models of biomolecular dynamics from HS-AFM imaging, thus overcoming the limitations of previous rigid-body fitting methods.

The chosen application of flexible fitting to HS-AFM data of the rather small HECT domain was challenging because the conformational dynamics observed under the AFM microscope is represented by topographies with a spatial resolution limited to a few tenths of pixels in each x and y scanning direction. While this limited information of tip-convoluted topographies should generally restrict the confidence of predictions from any fitting procedure, our results obtained from flexible fitting are very much consistent with previous structural data analysis and molecular modelling emphasizing the importance of flexible C-lobe domain rotations for HECT function. HECT dynamics observed in HS-AFM experiments corresponds to transitions between the spherical and L-shape protein shapes under equilibrium conditions which may regulate binding of the ubiquitin delivering E2 enzyme, known to preferably bind to the HECT domain in its L-shape conformation [21]. The role of flexible motions of the finger loop in the HECT N-lobe region is so far much less explored (to the best of our knowledge), but its proximity to the E2 enzyme binding site may suggest involvement in formation of the HECT-E2 complex. Further HS-AFM experiments considering HECT mutants are planned to investigate this aspect at the single molecule level. A very recent experimental work has indeed characterized HECT domain C-lobe and N-terminal region as key structural elements for ubiquitination in the HECT-type E3 ligase Ufd4 [24].

We expect the flexible fitting method to generally perform better in applications to HS-AFM imaging of larger proteins, where domain motions are visualized by kilopixel resolution (and beyond) topographies. Apparently, high quality imaging data obtained with a sharp scanning tip are favorable to improve the quality of atomistic model predictions.

The application of flexible fitting to the actin filament demonstrates scalability of the method, allowing for the atomistic-level interpretation of HS-AFM imaging far beyond the range of single protein topographies at reasonably low computational cost compared to the surreal expense required by molecular dynamics. Within the NMFF-AFM method, the generation of feasible dynamic conformations of a biomolecular structure is generally fast, but the computational efficiency of the entire fitting process is mainly gained from parallelized GPU calculations of simulation AFM images within the BioAFMviewer. All applications of flexible fitting were performed on standard laptop computers, with computations taking between tens of seconds to less than a minute for the analysis of a single AFM frame (except for the actin filament case).

The new software implementation of the NMFF-AFM method therefore opens the opportunity for large-scale analysis of scanning probe microscopy imaging data. The presented application for reconstructing an atomistic molecular movie of HECT protein dynamics from a sequence of fifty HS-AFM movie frames demonstrates a first advancement in that direction.

In principle, the computationally efficient application of flexible fitting can be extended to the automated analysis of *all* measured topographies and their dynamic changes accumulated from the time course of experimental observations. While this ambitious aim was not attempted in this work, we nonetheless discuss several challenging tasks currently preventing fully automated large-scale application of flexible fitting. The implemented method of automatized molecule recognition from HS-AFM imaging data can reliably provide individual target topographies for the presented flexible fitting applications, but it would generally not be possible to detect a single molecule topography in AFM images of clustered molecules or distinguish between different molecule types.

Further challenges for automation are related to the flexible fitting method. As demonstrated before in applications to synthetic AFM data, the performance of flexible fitting, and therefore the reliability of the obtained dynamic model conformation, may be affected by the chosen molecular orientation of the initial structure [15]. Within the BioAFMviewer workflow, the rigid-body fitting app provides an automatized procedure for initial alignment of the starting structure with respect to the target AFM image prior to the application of flexible fitting. However, for the application to the HECT domain the predicted alignments were not reliable because simulated AFM images of the starting structure are very different from the target HS-AFM image. In that case, our choice for the initial alignment was based on information about the molecule placement relative to the AFM substrate acquired from previous applications of rigid-body fitting of the starting structure to a selected set of HS-AFM images that represented HECT protein topographies close to the same equilibrium conformation [19]. We argue that this application of flexible fitting is an exemplary case where stark differences in the topography shapes of initial and target image may present a general challenge for confident initial molecule alignment by automation. Nonetheless, manual intervention to determine the initial alignment allowed us to further reconstruct an atomistic-precision molecular movie of dynamic models, including motions of large-amplitude conformational transitions, inferred by the completely automatized application of flexible fitting to fifty measured HS-AFM imaging frames.

The presented application to the HS-AFM image of HECT-E2 complex provides a demonstration of the ideal BioAFMviewer fitting workflow, consisting in reliable automated rigid-body alignment and the estimation of optimal tip-shape parameters prior to the actual flexible fitting process.

In summary, with the prospects of flexible fitting for large-scale automated analysis of AFM imaging data, the provided NMFF-AFM software implementation can be viewed as a first advancement towards that direction. Along this path, with several future challenges ahead, the presented contribution allows for the immediate practical application of flexible fitting to AFM measurement data within the versatile user-friendly BioAFMviewer software interface. The BioAFMviewer software is available as a free download from the project website www.bioafmviewer.com. All new presented developments are implemented in the BioAFMviewer version 5.0.

## Methods

### Implementation of the BioAFM-ASD viewer

We have developed our own computer code to process AFM experimental raw data encoded in the ASD file format and visualize measured topographic images. The ASD viewer app includes post-experimental image processing for AFM-stage tilt correction and for Gaussian filtering of AFM pixel information along the scanning grid. For the implementation we employed previously described algorithms [22]. For the automated detection of measured topographies to be considered as target AFM images, we calculated for each ASD frame the probability density of topography heights and then applied the Otsu method [20] to determine the height threshold separating relevant information from noisy background (the opencv library was used). The boundary for each isolated topography in the actual ASD frame was then characterized by a box with Nx pixels width and Ny pixels height. The corresponding region of interest (ROI) was constructed by extending the box size by specified pixel units. Alternatively, the ROI can be manually drawn within the ASD frame pixel canvas. The size of the target AFM image is fixed by the ROI, but there is an option to proceed fitting with either all topography height information retained, or the denoised case from application of the Otsu method threshold.

### Communication between the ASD viewer and BioAFMviewer simulation AFM

Once the fitting toolbox is activated, the scan step to generate simulated images is fixed to the value extracted from the loaded experimental ASD file. The ROI for a molecule in the ASD canvas would result in the same-size ROI centered at the origin in the simulation AFM canvas. In both image canvases, the ROIs can be moved but not resized as they must have the same size to be compared by the image similarity scoring functions.

### Implementation of the flexible fitting toolbox

The flexible fitting toolbox app integrates all computations of the NMFF-AFM iteration scheme and allows users to modify parameters of the model and the fitting process.

The only model parameter is the elastic network cutoff distance.

#### Elastic network cutoff distance

the cutoff distance determines the connectivity of atoms within the elastic network constructed from the input PDB structure, i.e. the pattern of effective harmonic interactions of each atom with neighboring atoms. The default value is chosen as 8 Å. Generally, a large cutoff distance would result in stiff elastic networks preventing normal modes to capture intramolecular flexibility. If the value is too small the elastic network would contain sparsely connected regions which give rise to unnatural motions (indicating the presence of additional trivial normal modes). In the latter case, the app will announce a warning alert.

Integrating the NMFF-AFM iteration scheme relies on the computation of normal modes of the instantaneous atomistic structure and the corresponding simulated AFM images of deformed conformations.

#### Computation of elastic network normal modes

To integrate the NMFF-AFM procedure into the BioAFMviewer, the original source code (see repository https://github.com/TamaLab/nmff-afm/tree/main) was rewritten in C++. For the normal mode analysis, the original Fortran source code (see repository https://github.com/TamaLab/nma) was compiled as a C++ library that was called within the BioAFMviewer iterative fitting procedure. Normal mode analysis is performed using the RTB method for computational efficiency [25]. The assignment of amino acid residues into blocks (residues per block, rpb) depended on the total number of atoms N of the biomolecular structure in the following way: N between 0-1000, 1 rpb; N between 1000-4000, 3 rpb; N between 4000-8000, 5 rpb; N between 8000-11000, 7 rpb; N > 11000, 11 rpb.

#### Simulation of AFM images

AFM simulation to translate atomistic structures into pseudo AFM topographies is at the core of the BioAFMviewer software. Simulated scanning is based on the non-elastic collisions of a rigid cone-shaped tip model (probe sphere radius *R*, and cone half angle ϑ) with the rigid van der Waals atomic model of the biomolecular structure (for details, see [17]). The calculation speed is massively enhanced by our implemented GPU-based atom filtering algorithm and computations [26]. Furthermore, the computation of simulated AFM images for all the deformed biomolecular conformations generated in each single iteration step is performed in parallel threads on the graphics card.

The following parameters are relevant for the fitting process.

#### Number of normal modes

Starting with the elastic network constructed from the initial input PDB structure the conformation is deformed in each iterative step according to a single favorable normal mode which is automatically selected from the number of specified normal modes. The total number of normal modes for an (anisotropic) elastic network of N atoms is 3N. The first 6 modes describe trivial translations and rotations of the structure and are required to include rigid-body rearrangements during iterative fitting. The remaining non-trivial modes 7…3N capture internal structural changes. Generally, first few non-trivial modes describe collective domain motions, whereas higher number modes correspond to more localized structural changes. By default, the number of modes is 15. Choosing a larger number of modes will increase the space of sampled conformational motions at the cost of additional computational time to complete fitting.

#### RMSD of structural deformation

The RMSD value quantifies the conformational change in each iterative step. It refers to the root mean square deviation (RMSD) of the atomic structure before and after the iteration step, i.e., 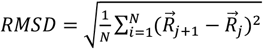, where *N* is the number of atoms. The 3N-dimensional vectors 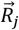 and 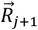 contain atomic positions of the structure before and after an iteration step *j*, respectively. Since 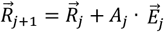, and normal mode vectors are normalized, the amplitude *A*_*j*_ chosen to apply a structure change according to the selected normal mode 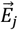 is given by the relation 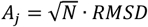. The default RMSD value is 0.5 Å. Using much larger values can result in unnatural structure changes, and it is recommended to use a larger number of iteration steps instead.

#### Image similarity scoring functions

To quantify the similarity between simulated AFM images of all deformed structures generated in each iteration step with the target AFM image we use scoring functions previously applied in computational AFM analysis. They are based on pixel-wise comparison between the simulated and measured height topographies across the scanning grid. The image correlation coefficient (image c.c.) is computed as 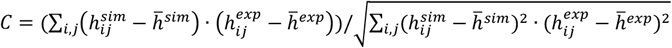. The cosine similarity (cos. sim.) considers absolute height values neglecting averages, i.e.,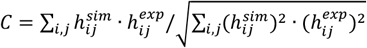. The pixel RMSD measures the root mean square deviation of height differences, i.e.,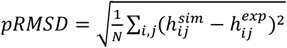. In the equations above, *h*_*ij*_ represents the height value for the pixel at position (*i, j*) of the scanning grid, and 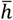 denotes the average of all pixel heights. The structural similarity metric (SSIM) is defined by 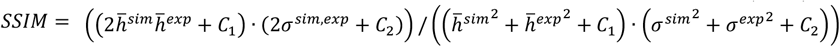, where *σ*^*sim*^ and *σ*^*exp*^ are the standard deviations of heights for the simulated and the experimental image, respectively, and *σ*^*sim,exp*^ is the covariance. The two constants *C*_1_ = (*k*_1_*L*)^2^ and *C*_2_ = (*k*_2_*L*)^2^ prevent divergence of the score. The value *L* represents the dynamic range of input data and is set as the maximum height found in the simulated and experimental image, and standard values *k*_1_ = 0.01 and *k*_2_ = 0.03 are used.

#### Number of iterations and criterion for automated termination of fitting

Flexible fitting terminates after a preset number of iterations. The default value is 100 steps, but the choice depends on the particular application. Additionally, we implemented a method for automated selection of a sufficiently fitted deformed structure based on *a posteriori* analysis of all iterations. Here, we follow the strategy formulated in the original paper [15] aimed to prevent structural deformations that only marginally increase image similarity of simulated and target experimental AFM image. This situation typically occurs when the image similarity is already sufficiently high, but fitting may continue to generate internal motions which cannot be justified due to the limited information provided by AFM surface topography imaging. To automatically select the sufficiently fitted structure, we traced the change of the image similarity score in each iteration, applied exponential fitting to the resulting data, and detected the iteration at which fitting should terminate by the criteria that it improved the score by a value less than 3-5 percent compared to the change at the initial iteration.

#### Fitting mode

We provide a choice between single-frame and multi-frame flexible fitting. In the first case, a single HS-AFM movie frame can be selected in the ASD viewer and fitting proceeds for a chosen topography (automatically detected or manually selected). For multi-frame fitting, aimed at reproducing a molecular movie from a time series of HS-AFM images, the user can specify the range of movie frames in the ASD viewer and individual fitting processes are performed frame by frame employing automated molecule detection to identify the corresponding target topography.

### Data analysis and graphical output

#### Structural change during fitting

The structural change during fitting is quantified by the root mean square displacement (RMSD) of atomic positions of the instantaneous conformation with respect to the initial conformation. The RMSD is also evaluated by alignment of the instantaneous conformation with the initial conformation, quantifying only internal conformational changes. Corresponding plots can be visualized after fitting and exported as image and text files.

#### Image similarity scores during fitting

The selected score, quantifying image similarity between the simulated and target AFM image along the fitting process, can be visualized as a line plot. Plots of the other available scoring functions are also provided for comparison, but it should be noted that fitting results employing them would be different. The data can be exported as image and text files.

#### Selected normal modes during fitting

Along the fitting process, the number of the normal mode selected in each iteration step is recorded and can be displayed as a point plot. The data can be exported as image and text files.

#### Error Map

The error map provides a graphical representation of differences between the simulated and target AFM topography. For the region of interest (ROI) the pair-wise comparison of height values for pixels across the scanning grid is performed and deviations are visualized by a color gradient. For the presented applications, deviations relative to mean heights were considered, i.e., for each cell of the scanning grid the height difference 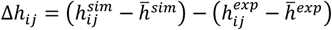 was evaluated. The visualization of the error map can be modified by color bar adjustments. The obtained data can be exported as images and text files.

#### Movie export

The molecular viewer and synchronized simulation AFM in the BioAFMviewer allow visualization of the fitting process as a molecular movie of atomistic motions from the initial orientation of the structure towards the conformation reached by fitting, together with the corresponding simulation AFM movie. In the case of the multi-frame flexible fitting, the atomistic molecular movie is displayed as a collection of predicted models for each frame together with the corresponding simulation AFM movie. Two types of visualizations are provided for export in the mp4 video file format. A side-by-side view of the molecular movie, the simulation AFM movie, and the target AFM image (movie). And second, a superposition of the molecular movie with the target AFM image (movie). Additionally, the molecular movie data can be exported as a multi-frame PDB file, and the simulation AFM movie data as an asd file, respectively.

### Details for application to HS-AFM imaging data

#### HECT domain

Flexible fitting was performed with the following set of parameters. Elastic network cutoff distance: 8 Å. Number of normal modes: 10. RMSD of structural deformation: 0.5 Å. Number of iterations: 100. Image similarity scoring: image c.c. The tip shape parameters for simulation AFM were: probe sphere radius R=2nm and cone half angle ϑ = 10° (Fig. 2, SI Movie 1), and R=1.0, 3.0 nm (SI Fig. 1, SI Movie 3).

#### HECT-E2 protein complex

Rigid-body fitting of the HECT-E2 protein complex was performed by sampling 3D molecular orientations with an angular grid spacing of 10° and using tip shape parameters R=2nm and cone half angle ϑ = 10° for simulation AFM. Application of the *Quick-Fit* procedure proceeded with the best matching molecular orientation based on image c.c. scoring, considering variations in the tip shape parameters (Δ*R* = ±1.5nm, step size 0.1nm; Δϑ = ±5°, step size 1°). The molecular orientation obtained with the optimal tip shape (R=2.3nm, ϑ = 14°) was used as the initial conformation for flexible fitting, which was performed with the following set of parameters: Elastic network cutoff distance: 8 Å. Number of normal modes: 15. RMSD of structural deformation: 0.5 Å. Number of iterations: 100. Image similarity scoring: image c.c.

#### HECT protein molecular movie

The experimental raw data was processed in the ASD-viewer to track the measured HECT topographies and determine the ROI for fitting within each frame. Flexible fitting was performed with the following set of parameters. Elastic network cutoff distance: 8 Å. Number of normal modes: 10. RMSD of structural deformation: 0.5 Å. Number of iterations: 100. Image similarity scoring: image c.c.

### HECT domain PDB structures for fitting and analysis

For the HECT domain application of flexible fitting the crystal structure PDB ID 3jvz (E6AP-type HECT catalytic conformation; chain C: Glu574-Gly950) was used. Other HECT domain structures PDB ID 1d5f (NEDD4L-type HECT L-shape conformation; chain A: Asn497-Ala846) and PDB ID 1nd7 (reverse T-shape; chain A: His544-Glu917) were considered for comparison of flexible fitting results with static structural data. The domain geometry of static HECT structures and dynamic conformations obtained from flexible fitting was characterized by the distance between C-lobe domain and N-lobe domain geometric centers and a hinge angle. For 3jvz the alpha-carbon atoms of residues Val838-Gly950 (C-lobe), Glu574-Asn746 and Asn783-Gly834 (N-lobe), and Asp747-Glu782 (finger-loop) were considered. For 1d5f the alpha-carbon atoms of residues Leu742-Ala846 (C-lobe), Asn497-Tyr645 and Glu685-Cys737 (N-lobe), and Glu646-Asn684 (finger loop) were considered. The orientation of the N-lobe domain in the respective HECT conformation was determined by computing the three principal axes of inertia. The hinge angle characterizing the orientation of the C-lobe relative to the N-lobe domain was then defined by the vector connecting the geometric centers and the major principal axis.

For the HECT-E2 application of flexible fitting, the crystal structure PDB ID 1c4z (HECT domain, chain A: Asn497-Ala846; E2 enzyme, chain D: Ser4-Tyr147) was used.

#### Actin filament

The 96 protomer atomistic model for fitting was constructed using the open source code provided in reference [27]. The resulting structure was already aligned along the x-axis in the coordinate system of our molecular viewer. Considering that actin bending towards the shape observed in the target HS-AFM image proceeds with the filament placed on the 2D substrate, allowed us to determine the degree of filament rotation around the x-axis required to prepare a suitable initial orientation for fitting. In other words, since global bending of the actin filament is described by motions corresponding to the first non-trivial normal mode (mode number 7), we have roughly aligned the filament such that bending motions would proceed within the x-y plane which represents the AFM substrate plane in our coordinate system. After that, we have additionally shifted the filament position in the x-direction such that the periodic height protrusion pattern in the simulated AFM image (coming from the actin helical pitch) roughly matches with that of the AFM image in the straight central region of the filament. It should be noted that a precise alignment is not necessary because further adjustments are taken care of during the flexible fitting process. To estimate the optimal tip shape for fitting, we compared cross-section height profiles of simulated AFM topographies for the initial orientation of the actin model with the measured HS-AFM topography. Using the BioAFMviewer topography analysis toolbox, we chose a cross section in the region where the filament is straight and varied the tip probe sphere radius value for computing simulated AFM images to roughly match the height profile in the measured image (see Fig. S4).

Flexible fitting was performed with the following set of parameters: Elastic network cutoff distance: 8 Å. Number of normal modes: 50. RMSD of structural deformation: 5 Å. Number of iterations: 100. Image similarity scoring: image c.c.

### HS-AFM experimental imaging

For the application of flexible fitting to HECT/E2 protein dynamics, we used a data set of previously obtained HS-AFM imaging and refer to the original publications [19] for further experimental details.

For the application of flexible fitting to the actin filament, original HS-AFM data was used. Experiments for actin were performed using a laboratory-built machine in tapping mode with small cantilevers (BLAC10DS-A2, Olympus) as previously described [28]. A DPPC:DPTAP:Biotin-Cap-DPPE (89:10:1) lipid bilayer was deposited (10 min) on the mica surface before the incubation of actin (10 min) (see Ref. [29] for detailed method of mica-supported lipid bilayer). A conventional actin buffer (25 mM KCl, 0.5 mM EGTA, 2 mM MgCl2, 20 mM Imidazol-HCl pH 7.0) was used for imaging.

### Computation benchmark details

All flexible fitting computations were performed on standard laptop computers. The computational efficiency evaluated using two architectures is summarized in Table 1.

**Table 1.** Computation benchmark details. Details of the two laptop computer architectures used for flexible fitting are provided together with the corresponding computation times for the four applications to HS-AFM imaging data.

|  | Single<br>HECT<br>image | HECT-E2 | HECT<br>movie<br>(50 images) | Actin |
| --- | --- | --- | --- | --- |
| **Lenovo ThinkPad X1 Carbon (7<sup>th</sup> Gen, 2019)**<br>CPU: Intel® Core™ i5-7200U 2.50GHz (up to 2.71GHz), 2 cores, 4 threads<br>RAM: 8GB<br>GPU: Intel® HD Graphics 620 | 29 sec | 44 sec | 26.5 min | 4.1 h |
| **Lenovo ThinkPad P14s (4<sup>th</sup> Gen, 2023)**<br>CPU: Intel® Core™ i7-1370P 1.90GHz (up to 5.20GHz), 14 cores, 20 threads<br>RAM: 64GB<br>GPU: NVIDIA RTX A500 Laptop, 4GB | 15 sec | 17 sec | 14 min | 1.5 h |

## Author contributions

Conceptualization: RA, OM, FT, HF.

Methodology: RA, OM, XW, FT, HF.

Software: RA, OM, FT.

Validation: RA, KT, NK, HK, OM, FT, HF.

Formal analysis: RA, HF.

Investigation: RA, KT, HK, NK, HF.

Resources: RA, HK, NK, OM, FT, HF.

Data curation: RA, NK, HK.

Writing – original draft: HF.

Writing – review & editing: RA, NK, HK, OM, FT, HF.

Visualization: RA, HF.

Supervision: OM, FT, HF.

Project administration: OM, FT, HF.

Funding acquisition: RA, OM, FT, HF.

## Acknowledgments

This work was supported by the Ministry of Education, Culture, Sports, Science and Technology (MEXT), Japan, through the World Premier International Research Center (WPI) Initiative (RA, NK, HK, and HF), Grant-in-Aid for Scientific Research KAKENHI Nos 25K01631 (HK) and 24H02260 (OM, FT) from the Japan Society for the Promotion of Science (JSPS), the Transdisciplinary Research Promotions grant (RA, NK, NK, and HF) from WPI-NanoLSI, and in part by MEXT as “Program for Promoting Researches on the Supercomputer Fugaku” (Development and application of large-scale simulation-based inferences for biomolecules JPMXP1020230119) (OM, FT).

## Supplementary Video captions (V1-V6)

**Video V1: Flexible fitting for HECT domain**. Visualization of large-amplitude conformation transitions during the fitting process in a side-by-side view of the molecular dynamics in the VdW representation of the protein domain, the corresponding sequence of simulation AFM images, the target HS-AFM image, and a superposition of the molecular dynamics with the target image.

**Video V2: Flexible fitting for HECT domain with denoised HS-AFM target image**. Visualization of large-amplitude conformation transitions during the fitting process in a side-by-side view of the molecular dynamics in the VdW representation of the protein domain, the corresponding sequence of simulation AFM images, the target HS-AFM image obtained after denoising, and a superposition of the molecular dynamics with the denoised target image.

**Video V3: Flexible fitting for HECT domain using different tip-shapes**. Visualization of large-amplitude conformation transitions during the fitting process in a side-by-side view of the molecular dynamics in the VdW representation of the protein domain, the corresponding sequence of simulation AFM images, the target HS-AFM image, and a superposition of the molecular dynamics with the target image. Top row: Flexible fitting results obtained with tip shape parameters (*R* = 3*nm*, ϑ = 10°). Bottom row: Flexible fitting results obtained with tip shape parameters (*R* = 1*nm*, ϑ = 10°).

**Video V4: Flexible fitting for HECT-E2 protein complex**. Visualization of E2 domain large-amplitude motions within the HECT-E2 complex during the fitting process in a side-by-side view of the molecular dynamics in the VdW representation of the protein complex, the corresponding sequence of simulation AFM images, the target HS-AFM image, and a superposition of the molecular dynamics with the target image.

**Video V5: Flexible fitting for HECT dynamics HS-AFM movie**. Side-by-side view of the atomistic molecular movie of HECT domain dynamics (in the VdW representation) obtained from multi-frame flexible fitting, the sequence of corresponding simulation AFM images, the sequence of target HS-AFM images, and a superposition of the molecular dynamics with the HS-AFM movie frames.

**Video V6: Flexible fitting for 4 megadalton actin filament**. Visualization of the fitting process showing the molecular dynamics in the VdW representation (top), the corresponding sequence of simulation AFM images (middle), and a superposition of molecular dynamics with the target HS-AFM image (bottom).

## Supplementary Figures (S1-S4)

**Fig. S1.**
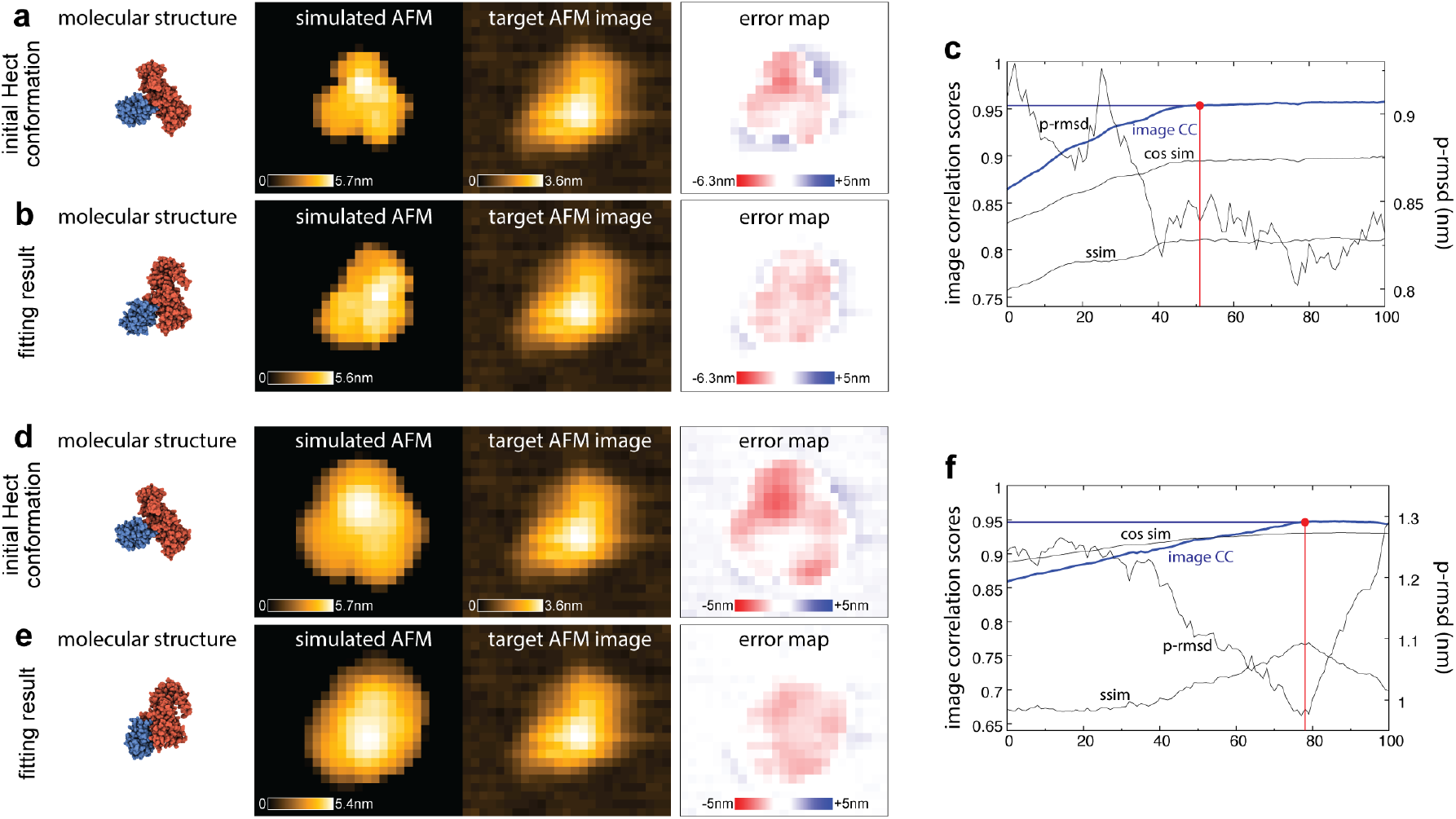
HECT domain large-amplitude conformational transition – fitting with different tip shape. **(a-c)** Flexible fitting results obtained with tip shape parameters (*R* = 1*nm*, ϑ = 10°). **(a)** Side-by-side image presentation of the initial HECT conformation for fitting (atomistic VdW representation), the corresponding simulated AFM image, the ROI of the target AFM image, and the error map. **(b)** The corresponding images after application of flexible fitting. **(c)** Plots of image similarity scores along the fitting process. The graph of the image cc score used for fitting is shown in blue; graphs of other scores are displayed in grey (for the p-rmsd an alternative scale is used). The iteration point determined by the fitting termination criterion is marked as a red dot. **(d-f)** Flexible fitting results obtained with tip shape parameters (*R* = 3*nm*, ϑ = 10°).

**Fig. S2.**
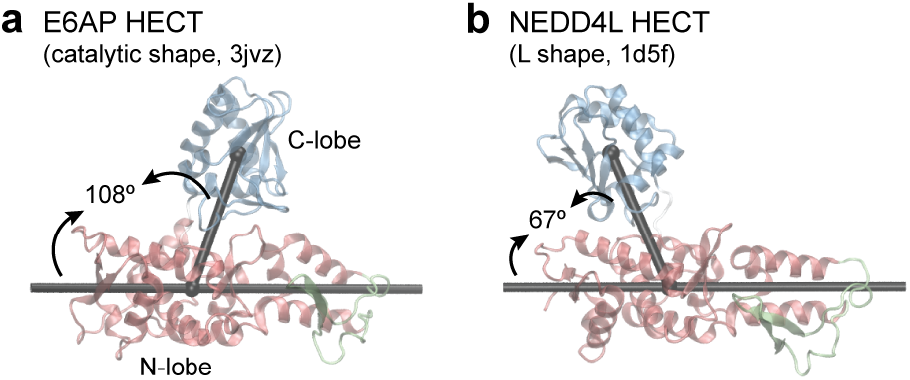
Geometry analysis of HECT crystal structures. **(a)** E6AP HECT domain in the catalytic shape state (PDB ID 3jvz) and **(b)** NEDD4L HECT domain in the L-shape (PDB ID d5f) shown in the ribbon representation. The C-lobe and N-lobe domains are colored in blue and red, respectively. Corresponding geometric centers are shown as beads in black color and their connection is indicated by a rod. The main principal axis of the N-lobe domain is also shown as a rod. For both structures the C-N-lobe hinge angle is indicated by arrows and corresponding values are given.

**Fig. S3.**
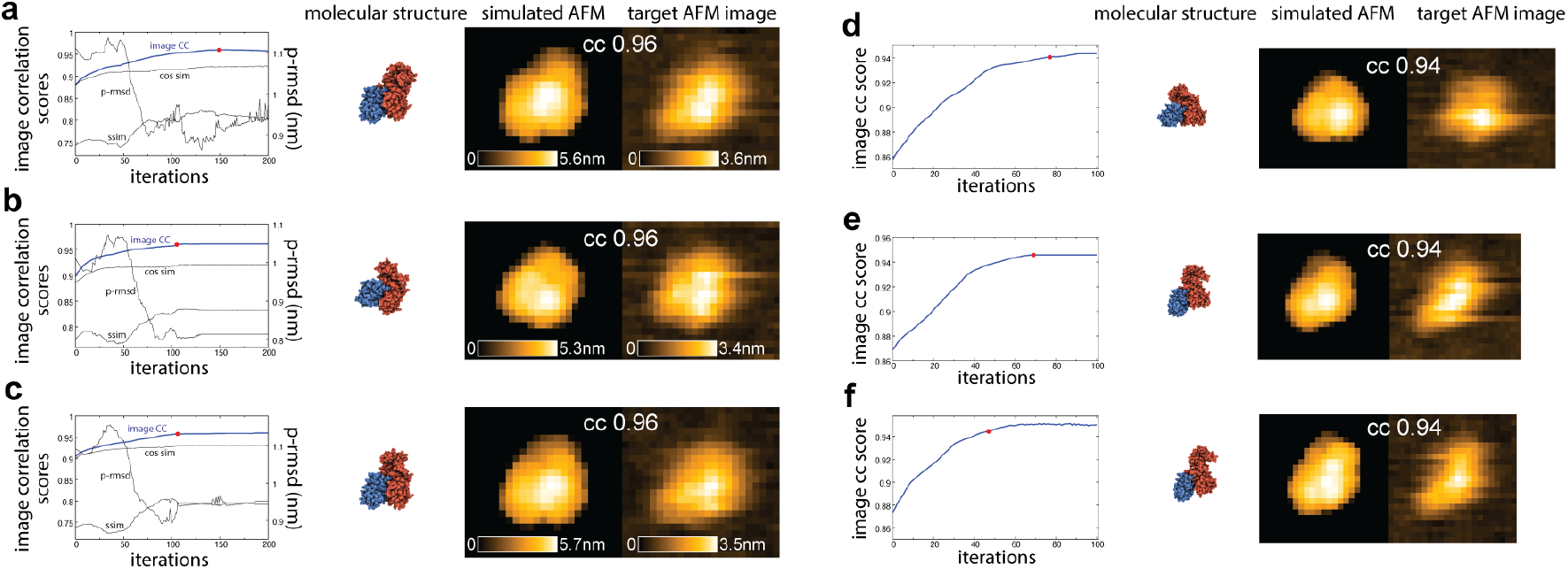
Reconstructing atomistic precision molecular movies from HS-AFM imaging. **Left side:** Flexible fitting results for three target experimental images from the HS-AFM movie set where fitting was extended to 200 iterations. In panels **(a-c)** plots of image similarity scores along the fitting process are shown on the left side. The graph of the image c.c. score used for fitting is shown in blue; graphs of other scores are displayed in grey (for the p-rmsd an alternative scale is used). The iteration point to terminate fitting is marked as a red dot. Improvement in fitting is quantified by a change in image cc values from 0.92 to 0.96 (a), 0.94 to 0.96 (b), and 0.94 to 0.96 (c). Shown on the right side are fitting results in a side-by-side image presentation of the HECT dynamic conformation (atomistic VdW representation), the corresponding simulated AFM image and the ROI of the target AFM image. **Right side:** Flexible fitting results for three target experimental images from the HS-AFM movie set whose measured topographies are much affected by artifacts.

**Fig. S4.**
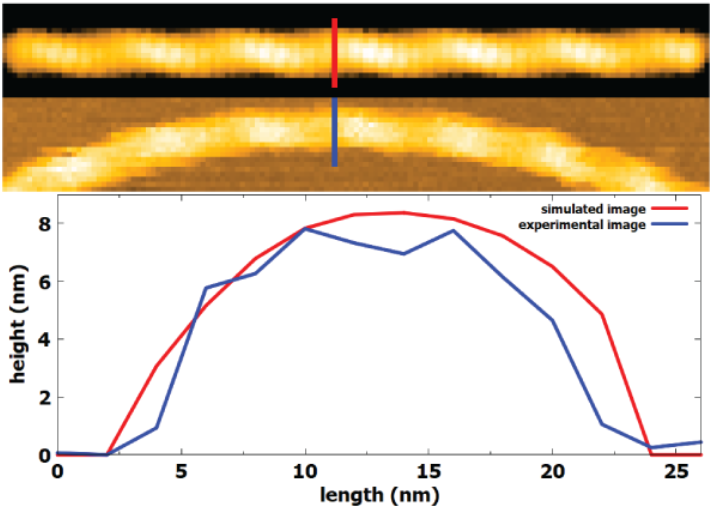
Tip shape estimate for actin filament flexible fitting. **Top:** Simulated AFM image of the atomistic actin model in its initial placement orientation obtained with tip shape parameters (*R* = 6.5*nm*, ϑ = 10°) together with the target HS-AFM image. **Bottom:** Topography heights monitored along the cross sections shown in red (simulated image) and blue color (experimental image), respectively.

**Methods Table 1**

